# Evaluation of antibacterial activity of ethanolic extracts of *Ruta chalepensis, Hippocratea excelsa, Litsea glaucescens* and *Waltheria americana* against methicillin-resistant *Staphylococcus aureus*

**DOI:** 10.1101/2024.09.27.615392

**Authors:** Margarita Salvador Martínez, José Fernando Fuentes Gutiérrez, Alonso Rubén Tescucano Alonso, Gabriel Martínez González

## Abstract

Antimicrobial resistance is still a problem today, with increasingly limited therapeutic options. *Staphylococcus aureus* causes infections of the skin, soft tissues and contains a high degree of pathogenicity. The objective of this work was to determine the antibacterial activity of ethanol extracts from several plants at different concentrations against strains of *Staphylococcus aureus* resistant and sensitive to methicillin, using the Kirby-Bauer method. The ethanolic extract of Rue (*Ruta chalepensis*) shows a significantly higher antibacterial activity than the extracts of Cancerina (*Hippocratea excelsa*), Laurel (*Litsea glaucescens*) and Tapacola (*Waltheria americana*) alone and in combination. Phenolic compounds, anthocyanins and flavonoids were determined. Rue may contribute to the potential identification of a natural alternative therapy against methicillin resistant *Staphylococcus aureus*.

## INTRODUCTION

*Staphylococcus aureus* (*S. aureus*) is a microorganism of great medical importance. For many years it has been recognized as one of the main human pathogens. *S. aureus* is part of the family *Microccocaceae*, genus *Staphylococcus*, which contains more than 30 different species and many of these are microbiota of the skin and mucous membranes in man. It is a Gram-positive coconut, not mobile. No spores can be found alone, in pairs, in short chains or clusters. It is an optional anaerobic but grows better in aerobic conditions. The microorganism produces catalase, coagulase and grows rapidly in blood agar. Their colonies are 1-3 mm long, produce a typical yellow pigment due to the presence of carotenoids and many strains produce hemolysis within 24-36 hours.^1^

*S. aureus* is a pathogen that can colonize mucous membranes and skin; and cause severe toxin-mediated invasive infections in humans and animals.^2^ Its spread has increased globally in adults and children, with clinical manifestations such as skin and soft tissue infections, more pathogenic pneumonia and bacteriemia associated with high morbidity and mortality rates, which has made them a public health problem.^3,4^

Currently, *S. aureus* strains have a wide range of antibiotic resistance and resistant and multi-resistant strains can be found. The acquisition of this resistance is mainly due to the horizontal exchange of genes that are carried by mobile genetic elements such as plasmids, transposons (Tn) and insertion sequences (IS).^1^ By the early 1960s, the genus *Staphylococcus* has developed resistance to many antibiotics that are available. Penicillin resistance of 80-93% or more is currently reported in isolated hospital and community strains of *S. aureus*. ^5,6^ Due to the resistance of *S. aureus* strains to penicillin, stable cephalosporins were introduced to semi-synthetic penicillases and penicillins in the late 1950s. This included methicillin, as the antibiotic of choice in the treatment of *S. aureus*. This drug was introduced to Europe in 1959 and a year later the first strain of *S. aureus* methicillin resistant (“methicillin resistant *S. aureus*”, SARM) was detected. Later, in 1963, the first nosocomial outbreak caused by MRSA strains was reported. Strains of *S. aureus* multiresistant have since been reported worldwide.^5-7^

Resistance is a factor that contributes to the development of complicated skin and soft tissue infections, limits the effect of some antimicrobial agents, what the development of new treatments requires.^8^ The use of natural products present in certain medicinal plants are fully suitable to prevent the growth of pathogens causing diseases, in particular some variants resistant to multiple drugs.^9^ The use of whole plants, bark, roots, leaves, etc., for the treatment of diseases such as respiratory complications has been a common practice in Africa and the Middle East for a long time.^10^ The aim of this study was to evaluate the antibacterial effect of ethanol extracts from Rue (*Ruta chalepensis*), Cancerina (*Hippocratea excelsa*), Laurel (*Litsea glaucescens*) and Tapacola (*Waltheria americana*) against *S. aureus* (SARM).

## MATERIALS AND METHODS

### Plant material and extraction

Rue (stem, flower, leaf), Tapacola (leaves and flower), Cancerina (bark) and Laurel (leaves) were used for the trials, purchased in the Sonora market of Mexico City. The plant material was identified taxonomically in the FES Iztacala-UNAM Herbarium, leaving a specimen that was integrated into the ethnobotanical collection of the herbarium. Each botanical material was macerated using 70% ethanol (HYCEL). It was left to rest for 15 days in the absence of light in a cool and dry place, making periodic movements. After time, the plant material was filtered and removed and the solvent evaporated at reduced pressure in the Rotaevaporator (Hahn Shin Scientific, HS-2000NS). The extract obtained was placed in an amber jar and stored at 4°C 2°C until use.

### Phytochemical screening

Colorimetric tests were performed to determine the presence of secondary metabolites present in the extracts [anthocyanins-HCl (MEYER); flavonoids-Shinoda method; phenolic compounds-ferric chloride reaction (MEYER)].^11^

### Antibacterial activity by disc diffusion method

Four strains, all of which were *S. aureus*, three of them methicillin-resistant, Ciprofloxacin and Erythromycin identified as 5, 10, 39^12^ and *S. aureus* ATCC 6538, were evaluated, taken from the ceparum of the Microbiology laboratory of the University of Ixtlahuaca CUI. Each strain was confirmed for purity (Gram stain), microbial identification (resealing in selective-differential and biochemical media) and was performed with four antibiotics [Cefoxitin 30 μg (BD BBL), Erythromycin 15 μg (BD BBL), Gentamicin 10 μg (BD BBL) and Ciprofloxacin 5 μg (BD BBL)].

From the macerated extract, different concentrations were prepared with Dimethylsulfoxide (DMSO) (EMSURE ACS), having the following: 1) for Rue, Cancerina and Tapacola were prepared at 25 mg/mL, 50 mg/mL, 100 mg/mL and 300 mg/mL); 2) for Laurel at 25 mg/mL, 50 mg/mL, 100 mg/mL and 500 mg/mL. The extracts were kept in refrigeration at 4 ºC.

Strains of *S. aureus* were planted by cross-strewn, in Soya Trypticaine (TSA) agar (DIBICO), incubating at 37°C for 24 hours, from the seed isolated colonies are taken, placing in tube with sterile saline solution, equalizing with standard tube 0.5 Mc Farland (1.5×10^8^ CFU/mL) and where the concentration was then corroborated by reading between 0.08 and 0.13 absorbance in the visible range spectrophotometer and UV 30% (Velab®) at a wavelength of 625 nm. Each standardized inoculum was sown by closed stria with the aid of a sterile swab in a Müeller-Hinton agar (DIBICO) box, then placed the AA discs with sterile tongs on each petri dish and 10 μL of the extract to be evaluated.

Each test was performed in triplicate. The results of antibacterial activity were analyzed using the SPSS statistical program-19, by means of an ANOVA test (Analysis of variance), comparison of Tukey averages (p ≤0.05).

Controls used: Negative control: Sterile disc with 10 μL of DMSO, sterile discs without anything, extracts impregnated in sterile discs [each control was placed in a tube with nutrient broth (DIBICO) and incubated at 37°C for 48 h]; Positive control: Linezolid antibiotic disc 30 μg (BD BBL).

## RESULTS AND DISCUSSION

Traditional Mexican medicine has played an important role in the treatment of various diseases. Plant products, particularly dry drugs and extracts have clearly moved from a dominant position as first-line treatment to disuse; However, in recent decades they have again reached a growing presence in medicine. This return has been facilitated by the search for alternatives to natural treatments such as the scientific development of plant medicines, seeking to reduce adverse effects, even if they are nil, in addition to the resistance generated in recent years and the increased knowledge of the risk-benefit of synthetic drugs.^13^

In this work we used medicinal plants, in particular Rue, Laurel, Cancerina and Tapacola (acquired in the market of Sonora of Mexico City), these are employed in different countries, Mexico being an entity of interest in these products for their multiple applications in herbal medicine, for example: Rue, it is attributed antiparasitic, cytotoxic, antiseptic and improvement of digestive problems and circulation, among others more^14^; Cancerina, is very popular in the states of Puebla, Morelos and Guerrero for its properties as antiseptic or curative in gastritis with infectious etiology and in cancer^15^. As for Laurel, it is an ornamental plant, being its common use in traditional cuisine as a condiment, also, its essential oil has great relevance in the cosmetic industry, mainly as flavouring^16^ and has been shown to have antibacterial activity against different microorganisms, including *S. aureus*^17^. Finally, the Tapacola plant has some therapeutic uses, in particular, digestive tract ailments and in cases of lesions and skin ulcers.

Taxonomic identification of the botanical material was carried out to know the grouping and classification that allowed to analyze the plant diversity in a rational and methodical way. At the same time, a specimen was left for collection in the FES Iztacala-UNAM herbarium (State of Mexico), which was assigned an internal registration number (see Table 1).

**Table 1.** Taxonomic identification of botanical material.

| Botanical material |  | Institution | Registration |
| --- | --- | --- | --- |
| Common name | Botanical name |  |  |
| Rue | *Ruta chalepensis* L. | Herbarium<br>FES- Iztacala<br>(UNAM) | 2673 |
| Cancerina | *Hippocratea excelsa*<br>Kunth |  | 2598 |
| Laurel | *Litsea glaucescens*<br>H.B.K |  | 2597 |
| Tapacola | *Waltheria americana* L. |  | 2674 |

Three strains of *S. aureus* resistant to methicillin (cefoxitine), ciprofloxacin, erythromycin and gentamicin were used, isolated in public health institutions^12^. Also, a strain of *S. aureus* susceptible to Methicillin. The current interest in studying this microorganism stems from its high frequency in clinical cases, which is worse, for presenting strains resistant to Methicillin. Consequently, one of the main causes of outbreaks of nosocomial infection in our country. However, it is not only a national problem but also extends to other countries. WHO^18^ created a list of three categories according to the urgency in which new antibiotics are needed, where *S. aureus* is within priority 2, classified as high urgency.

Since the search for alternative therapies was important, the antibacterial activity of the extracts obtained against different strains of *S. aureus* was evaluated, obtaining that the ethanolic extract of Rue (*Ruta chalepensis*) showed the greatest effect of the evaluated extracts, against all strains used, obtaining inhibition halos of 15.67 ± 1.38 mm up to 24.46 ± 0.24 mm at a concentration of 300 mg/mL, slightly below the control of Linezolid with an inhibition halo of 32.88 ± 0.76 mm, despite being under the control, allows us to define that it is having an action on resistant strains that have generated mechanisms to evade the action of antibiotics and would not check if the action observed is bactericidal or bacteriostatic (See Tables 2-5).

**Table 2.** Antibacterial activity of *S. aureus* strain No. 5 (SARM) against different plant extracts.

| Extract | Concentration | Average<br>(Inhibition halo<br>mm) ± Standard<br>deviation | p-value | |
| --- | --- | --- | --- | --- |
| Rue | 50 mg | $9.16 \pm 1.10^*$ | 300 mg vs<br>50 y 100mg | 0.000 |
| | 100 mg | $13.06 \pm 2.46^*$ | | 0.000 |
| | 300 mg | $21.08 \pm 0.84$ | | |
| Cancerina | 25 mg | $7.31 \pm 0.27^*$ | 300 mg vs 25,<br>50 y 100mg | 0.000 |
| | 50 mg | $6.44 \pm 0.42^*$ | | 0.000 |
| | 100 mg | $10.80 \pm 0.26^*$ | | 0.023 |
| | 300 mg | $12.55 \pm 0.98$ | | |
| Laurel | 25 mg | $7.96 \pm 0.27^*$ | 500 mg vs 25,<br>50 y 100mg | 0.000 |
| | 50 mg | $8.90 \pm 0.27^*$ | | 0.000 |
| | 100 mg | $10.16 \pm 0.25^*$ | | 0.002 |
|  | 500 mg | 12.19 ± 0.65 |  |  |
| Tapacola | 50 mg | 7.84 ± 0.74* | 300 mg vs 50 y 100mg | 0.000 |
|  | 100 mg | 10.44 ± 0.60* |  | 0.008 |
|  | 300 mg | 13.57 ± 1.48 |  |  |
| T/R | 300 mg/ 300 mg | 16.83 ± 0.86* | 300 mg Rue vs T/R | 0.008 |
| C/R | 300 mg/ 300 mg | 18.81 ± 0.53 | 300 mg Rue vs C/R | 0.277 |
| Combination R-C-L-T | 50 mg/25 mg/ 25mg/ 100 mg | 8.30 ± 0.22* | 300 mg Rue vs R-C-L-T | 0.000 |
| Control (positive) | Linezolid 30 µg | 32.12 ± 0.75* | 300 mg Rue vs Control | 0.000 |

\*T/R: Tapacola/Rue, C/R: Cancerina/Rue, Combination R-C-L-T: Rue/Cancerina/Laurel/Tapacola. SARM: *S. aureus* Resistant to Methicillin. \*Significant difference (TUKEY, $p \leq 0.05$ ).

**Table 3.**
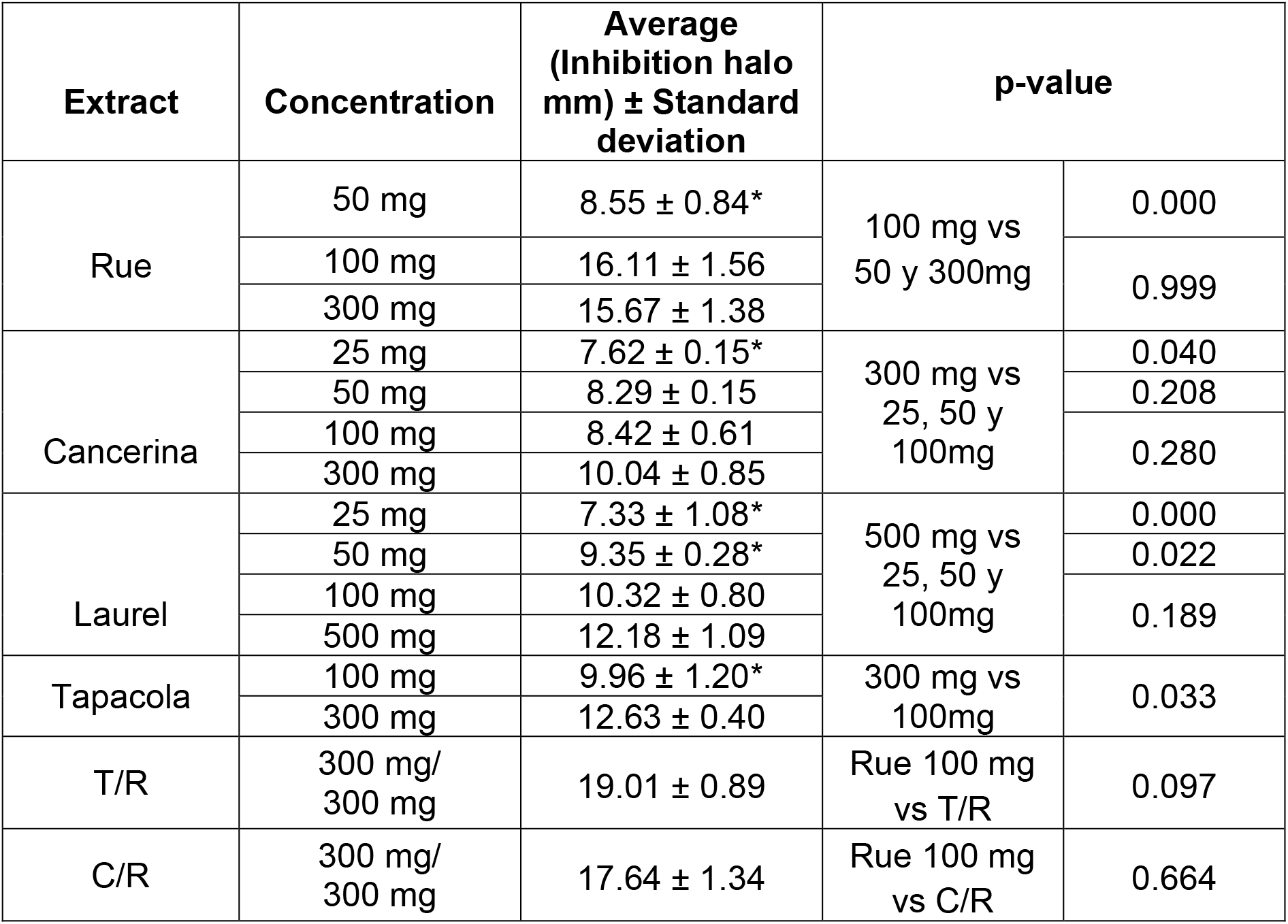
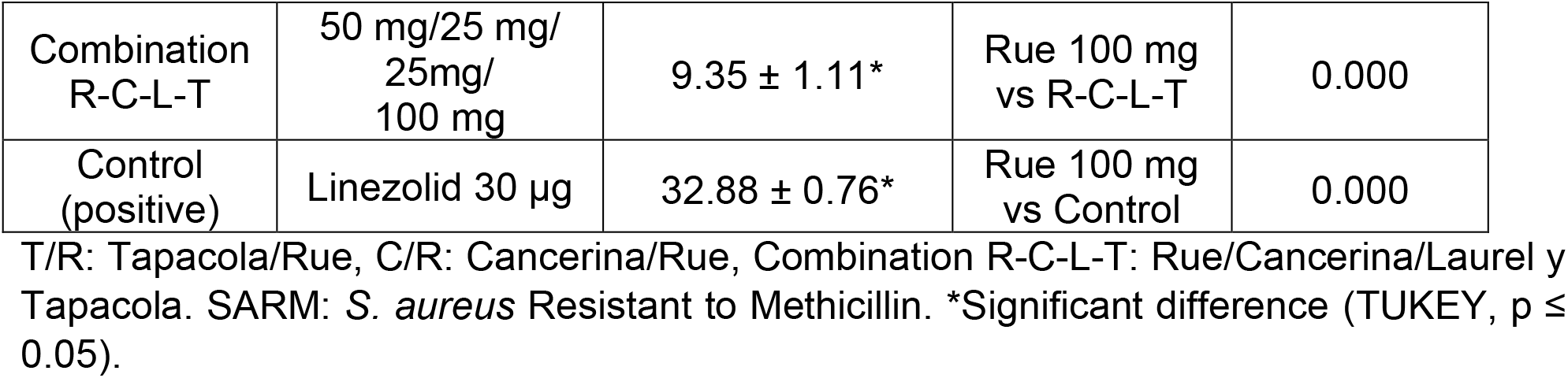
Antibacterial activity of *S. aureus* strain No. 10 (SARM) against different plant extracts.

In contrast, compared with other investigations, the concentrations and inhibition halos vary between authors of some reviews; Rodrigues et al.^19^ and Mohammed^20^ found no activity in the Rue extract (*Ruta chalepensis*) against *S. aureus*; while Ouerghemmi et al.^21^, evaluated activity in flowers, leaves and stem at a concentration of 5 mg/disc, finding inhibitions from 15 ± 0.6 mm to 16.3 ± 0.6 mm; Bonjar et al.^22^, evaluated antibacterial activity at a concentration of 20 mg/mL and obtained an inhibition halo average of 10 mm, same activity found Alzoreky et al.^23^, at a concentration lower than 10 mg/mL; Toribio et al.^24^ obtained a halo of 16 mm from 20 g methanol extraction of dried aerial parts, when compared with the results obtained in this work it is observed that a similar inhibition (9.92 ± 0.38 mm) is achieved, but at a higher concentration (50 mg/mL).

At the same time, the highest antibacterial effect was 24.46 ± 0.24 mm at a concentration of 300 mg/mL, like the study by Ivanova^25^, which reported an inhibition halo of 23 mm, but at a lower concentration (0.5 mg/mL).

As for the Cancerina extract (*Hippocratea excelsa*), it shows greater inhibition at a concentration of 300 mg/mL, the inhibition halos were 10.04 ± 0.85 mm up to 12.55 ± 0.65 mm, although there was very low or no inhibition at the minimum concentration of 25 mg/mL.

However, the inhibition halos of Laurel extract (*Litsea glaucescens*) corresponded to

7.33 ± 1.08 mm up to 12.86 ± 0.61 mm at a concentration of 500 mg/mL, there is an antibacterial effect, however, it is very little, in contrast to the study reported by Ouibrahim et al.^26^, although it differs from the extraction method, using an essential oil as final product and obtaining inhibition halos of 8.4 to 22.4 mm; however, Millezi et al.^17^, evaluated the Laurel extract against four microorganisms, including *S. aureus*, of all concentrations only had activity at the highest concentration of 50%, with an inhibition halo of 8 mm, much lower than that obtained in this research work.

In contrast, the inhibition halos obtained from the extract of Tapacola (Waltheria americana) were 10.64 ± 0.59 mm to 13.57 ± 1.48 mm at a concentration of 300 mg/mL, however, when comparing the results with what is reported, there is only information reporting antibacterial activity, but no studies supporting it, only the work done by Okwute et al.^27^, who work with a different species (*Waltheria indica*), where it obtained an inhibition halo of 23 ± 0.15 mm at a concentration of 10 mg/mL.

On the other hand, it was decided to make combinations between the extracts, to know if there is a synergistic effect that potentiated the inhibitory effect. Of all the possible combinations, it was observed that Rue, at a concentration of 300 mg/mL, presented the greatest inhibitory effect compared to the other extracts, it was combined with Tapacola and Cancerina, which were the extracts following a better antibacterial activity, both at a concentration of 300 mg/mL. Also, the possible toxicity that could present the components of plants was considered, so it was decided to handle minimal concentrations of the four extracts that had antibacterial activity being 50 mg, 100 mg, 25 mg and 25 mg: for Rue, Tapacola, Laurel and Cancerina respectively. Results of the 1:1 (300 mg/mL) combination of Rue with Tapacola, inhibition halos were obtained from 16.83 ± 0.86 mm to 21.43 ± 2.84 mm, when compared with individual results for Rue 24.46 ± 0.24 mm and Tapacola 13.57 ± 1.48 mm The inhibition halos were lower. Mainly, it is observed that there was a synergistic effect with respect to the extract of Tapacola alone, whereas the extract of Rue at a concentration of 300 mg/mL continues to have the greatest effect despite the combination (see Table 2-4).

**Table 4.** Actividad antibacteriana de la cepa No. 39 de *S. aureus* (SARM) contra diferentes extractos de plantas.

| Extract | Concentration | Average (Inhibition halo mm) ± Standard deviation | p-value |  |
| --- | --- | --- | --- | --- |
| Rue | 50 mg | 9.92 ± 0.38* | 300 mg vs 50 y 100mg | 0.000 |
|  | 100 mg | 16.21 ± 2.46* |  | 0.001 |
|  | 300 mg | 24.46 ± 0.24 |  |  |
| Cancerina | 25 mg | 6.96 ± 1.31* | 300 mg vs 25, 50 y 100mg | 0.004 |
|  | 50 mg | 7.27 ± 1.01* |  | 0.007 |
|  | 100 mg | 10.53 ± 1.37 |  | 0.975 |
|  | 300 mg | 11.28 ± 1.04 |  |  |
| Laurel | 25 mg | 9.53 ± 0.93 | 300 mg vs 25, 50 y 100mg | 0.295 |
|  | 50 mg | 9.55 ± 0.92 |  | 0.307 |
|  | 100 mg | 10.76 ± 0.16 |  | 0.569 |
|  | 500 mg | 11.35 ± 1.04 |  |  |
| Tapacola | 50 mg | 7.58 ± 0.66* | 300 mg vs 50 y 100mg | 0.004 |
|  | 100 mg | 8.08 ± 0.27* |  | 0.006 |
|  | 300 mg | 12.94 ± 0.25 |  |  |
| T/R | 300 mg/300 mg | 21.43 ± 2.84 | Rue 300 mg vs T/R | 0.405 |
| C/R | 300 mg/300 mg | 22.61 ± 1.22 | Rue 300 mg vs C/R | 0.854 |
| Combination R-C-L-T | 50 mg/25 mg/25mg/100 mg | 9.65 ± 0.39* | Rue 300 mg vs R-C-L-T | 0.000 |
| Control (positive) | Linezolid 30 µg | 32.00 ± 1.02* | Rue 300 mg vs Control | 0.002 |

T/R: Tapacola/Rue, C/R: Cancerina/Rue, Combination R-C-L-T: Rue/Cancerina/Laurel y Tapacola. SARM: *S. aureus* Resistant to Methicillin. \*Significant difference (TUKEY, $p \leq 0.05$ ).

**Table 5.**
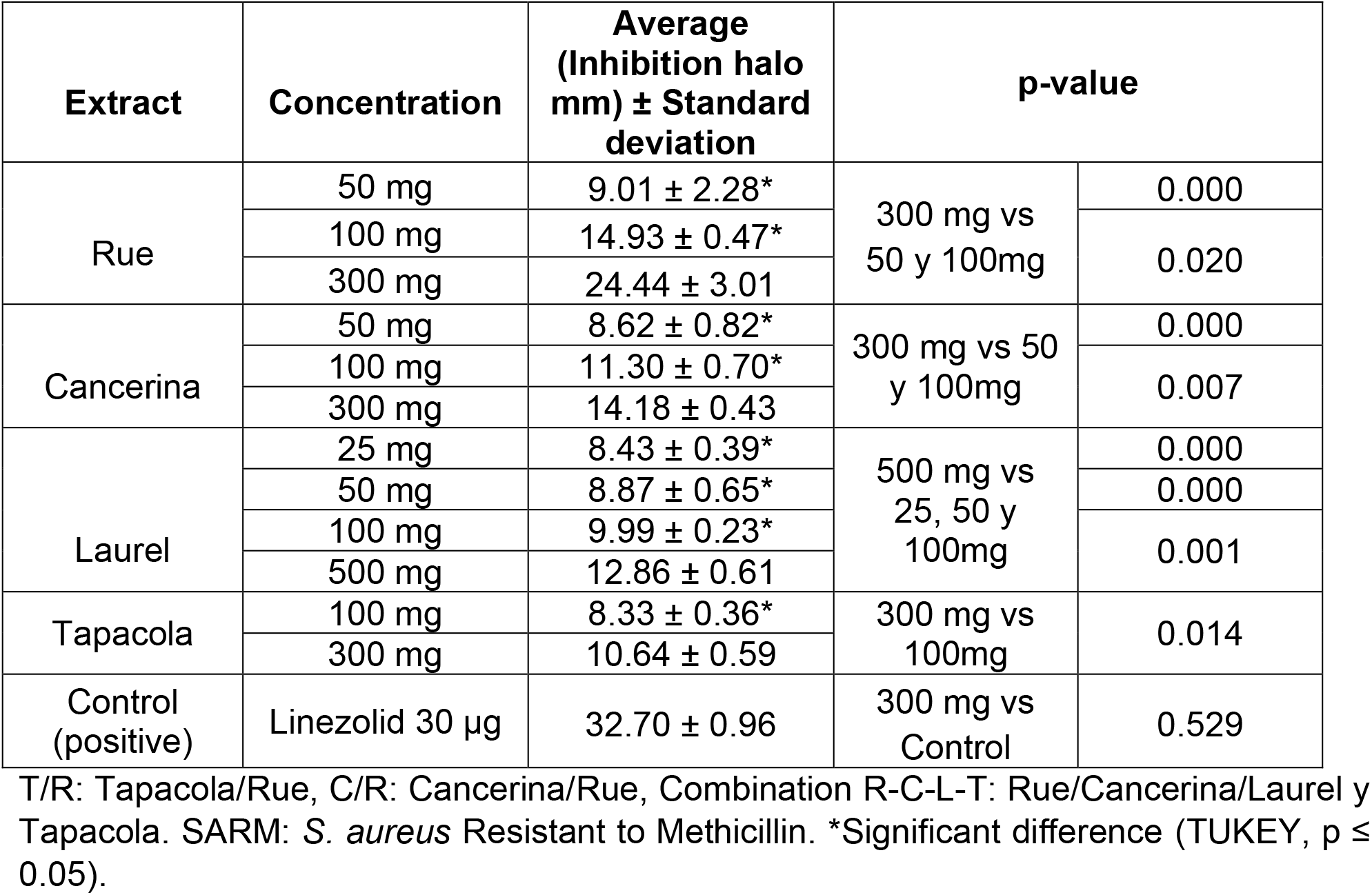
Antibacterial activity of the *S. aureus* strain ATCC 6538 against different plant extracts.

In relation to the combination of the four extracts at lower concentrations inhibition halos were achieved from 8.30 ± 0.22 mm to 9.65 ± 0.39 mm, that is, a decrease in inhibition can be observed, so that there is an antagonistic effect by some or several extracts being in combination, so that it is not recommended to use these extracts in combination as possible therapy against *S. aureus*.

As regards the results of antibacterial activity, they differ from several studies in other countries, partly due to agroclimatic conditions, plant age, type of species, type of plant material used (leaves, flowers, stems) that they were able to generate different compounds, certainly necessary for their development, adaptation and survival, which are a fundamental part of the antibacterial activity.

The presence of phenolic compounds, flavonoids and anthocyanins (see Table 6) were determined in the phytochemical tests on the extracts evaluated, possibly conferring the antibacterial effect. Also, the presence of phenolic compounds in Rue, coincide with what was reported by Naveda et al.^14^, where he performed a phytochemical march to stems, flowers and leaves, consequently positive to the test of phenolic compounds, perhaps, with antibacterial effect against *S. aureus*. Flavonoids are also derived from phenolic compounds, have in their chemical structure a variable number of phenolic hydroxyl groups, which easily penetrate the bacterial cell membrane, bind and precipitate protoplasmic proteins, denaturation, that is, they act as protoplasmic poisons.^28^ Another aspect, it is likely, that the location and number of hydroxyl groups in the phenol group are related to the toxicity of polyphenols against the microorganism.^29^

**Table 6.** Identification of secondary metabolites from extracts (Laurel, Cancerina, Rue and Tapacola).

| Test | Extract |  |  |  |
| --- | --- | --- | --- | --- |
|  | Laurel | Cancerina | Rue | Tapacola |
| **Anthocyanins** | + | + | + | + |
| **Flavonoids**<br>**(Shinoda)** | + | + | + | + |
| **Phenolic compounds**<br>**(iron chloride)** | + | + | + | + |

+: Presence (qualitative test).

Also, these flavonoids cause bacterial death by inhibiting the synthesis of ribonucleic acid or deoxyribonucleic acid, due to their flat structure similar to that of purine and pyrimidic bases, they can therefore intermingle forming hydrogen bonds with the bases in single or double chain and thus the flavones alter the three-dimensional structure of nucleic acids, preventing their proper de novo synthesis, as a result, cause reading errors during transcription.^30^

Finally, it is suggested to carry out more studies in Rue or Tapacola, since the extracts with best activity inhibited the growth of S. aureus strains resistant to Meticillin, In particular, the metabolites of plant extracts should be used as a therapeutic alternative in future.

## CONCLUSIONS

The four extracts from different plants had an inhibitory effect so that the hypothesis is finally proven. Rue extract shows the best antibacterial effect against Methicillin-resistant *Staphylococcus aureus* and Methicillin-sensitive *Staphylococcus aureus*. The extract of Rue (*Ruta chalepensis L*.) according to phytochemical tests contains phenolic compounds to which this antibacterial effect is attributed, due to its different known mechanisms of action.

The result of this research contributes to the potential determination of a natural alternative therapy against Methicillin-resistant *S. aureus* (MRSA).

## THANKS

To the University of Ixtlahuaca CUI, to the Q.B.P Olga Mateos Salazar, to the coordination of Q.F.B and to the Herbarium of the FES-Iztacala-UNAM, for support in carrying out this research work.

## «Conflicts of interest

none»

## Notes

### Competing Interest Statement

The authors have declared no competing interest.

